# Combined Enhanced Biological Phosphorus Removal (EBPR) and Nitrite Accumulation for Treating High-strength Wastewater

**DOI:** 10.1101/2021.01.16.426983

**Authors:** Zhihang Yuan, Da Kang, Guangyu Li, Jangho Lee, IL Han, Dongqi Wang, Ping Zheng, Matthew C. Reid, April Z. Gu

## Abstract

The enhanced biological phosphorus removal (EBPR) has been widely applied in treating domestic wastewater, while the performance on high-strength P wastewater is less investigated and the feasibility of coupling with short-cut nitrogen removal process remains unknown. This study first achieved the simultaneous high-efficient P removal and stable nitrite accumulation in one sequencing batch reactor for treating the synthetic digested manure wastewater. The average effluent P could be down to 0.8 ± 1.0 mg P/L and the P removal efficiency was 99.5 ± 0.8%. *Candidatus* Accumulibacter was the dominant polyphosphate accumulating organism (PAO) with the relative abundance of 14.2-33.1% in the reactor. Examination of the micro-diversity of *Candidatus* Accumulibacter using 16s rRNA gene-based oligotyping analysis revealed one unique Accumulibacter oligotype that different from the conventional system, which accounted for 64.2-87.9% of the total Accumulibacter abundance. The presence of high-abundant glycogen accumulating organisms (GAO) (15.6-40.3%, *Defluviicoccus* and *Candidatus* Competibacter) did not deteriorate the EBPR performance. Moreover, nitrite accumulation happened in the system with the effluent nitrite up to 20.4 ± 6.4 mg N/L and the nitrite accumulation ratio was nearly 100% maintained for 140 days (420 cycles). *Nitrosomonas* was the dominant ammonia-oxidizing bacteria with relative abundance of 0.3-2.4% while nitrite-oxidizing bacteria were almost undetected (<0.1%). The introduction of extended anaerobic phase and high volatile fatty acid concentrations were proposed to be the potential selector forces to promote partial nitrification. This is the first study that combined EBPR with nitrite-accumulation for digested manure wastewater treatment, and it provided new sights in strategies to combine the EBPR and short-cut nitrogen removal via nitrite to achieve simultaneous nitrogen and phosphorus removal.

## 1. Introduction

High-strength wastewaters refer to the wastewater containing higher concentration of contaminants than domestic wastewater which can be tracked from different sources such as sludge digestion (centrate), agricultural, textile, food, or other industries (Mutamim et al., 2012, Hamza et al., 2016). Agricultural wastes, such as manure wastes, belong to the high-strength waste streams that comprise relatively high concentration of soluble chemical oxygen demand (sCOD) as 700-2500 mg/L, ammonia as 47-1300 mg N/L and phosphate as 10-120 mg P/L (Liu et al., 2014). For wastewater treatment, biological technologies are being explored as the preferred options due to potential economic advantages and sustainability considerations. Currently, anaerobic digestion (AD) has been widely applied in treating the centrate, manure, and other industrial wastewaters containing high-strength organic matter to achieve energy recovery and lower the carbon footprint (Nasir et al., 2012). However, the AD process still leaves the nutrient (mainly nitrogen and phosphorus) behind, and high-strength nutrient, such as those in manure digester effluent discharge, could cause severe environmental problems such as eutrophication and threaten the ecosystem quality, water security, and public health (Dodds et al., 2009).

For nutrients removal and recovery from anaerobic digestion effluent, anaerobic ammonium oxidation (Anammox) has been promoted as a promising cost-efficient biological nitrogen removal technology, which has been successfully applied for treating high-strength ammonium-containing wastewaters (500-1500 mg N/L) around the world (Kuenen, 2008, Lackner et al., 2014). Although the successful implementation of anammox process for treating high-strength wastewater has been demonstrated, achieving the stable nitritation process to enable short-cut of the conventional nitrification process for relatively low-strength ammonium wastewater (30-100 mg N/L) remains challenging (Cao et al., 2017).

In contrast to the extensive attention on the innovative technologies targeting only at removing the high-strength ammonium in the wastewater, phosphorus removal and high-strength P-related issues have hardly been addressed. Enhanced biological phosphorus removal (EBPR) has been widely applied in treating domestic wastewater, however, the application of EBPR for high-strength waste streams has hardly been explored. EBPR has been deemed as unsuitable due to the presumptions that the high-strength wastewater characteristics may not be suitable for polyphosphate accumulating organisms (PAOs), as well as concerns regarding the performance stability that are susceptible to microbial community competitions and other factors such as pH, temperature, solids retention time, influent carbon to phosphorus ratio and system configurations (Neethling et al., 2006, Onnis-Hayden et al., 2020). A limited number of studies using lab-scale EBPR reactors to treat synthetic high-strength wastewater showed promising overall P removal efficiency (~70%), but no process optimization and comprehensive evaluations of kinetics and mechanisms of the EBPR were performed in these studies (Bickers et al., 2003).

Whether EBPR can be combined with short-cut N removal processes for high strength wastewater treatment remains largely unexplored. There are several recognized conflicts in combining EBPR with short-cut N removal processes. For example, EBPR requires sufficient carbon to phosphorus ratio (C/P) in the influent and short solids retention time (SRT), whereas the short-cut N removal process prefers low or no carbon and long SRT. Moreover, it was also presumed that the slow-growing bacteria such as autotrophic nitrifiers cannot compete well with the fast-growing heterotrophic bacteria including PAOs, resulting in the incompatibility of the N and P removal process in one reactor. For the short-cut N removal process, nitrite is desired to be the intermediate product rather than nitrate, and some denitrifying PAOs (DPAOs) can use nitrite as electron acceptor instead of nitrate to achieve N and P removal simultaneously (Jiang et al., 2006, Gao et al., 2019), whereas, the high ammonia and nitrite accumulation could inhibit the normal PAOs activity (Zheng et al., 2013, Jabari et al., 2016). Low dissolved oxygen (DO) was assumed to be an effective engineering control strategy to achieve nitritation process (Chuang et al., 2007, Brockmann and Morgenroth, 2010), but can also limit the P uptake rate of PAOs.

Recent research advances in fundamental understanding and successful full-scale implementation of influent carbon-independent, return activated sludge (RAS)-fermentation S2EBPR (side-stream enhanced biological phosphorus removal) process achieved concurrence of PAOs selection in presence of sludge digested centrate and promises new opportunities for its application for P-rich wastes (Onnis-Hayden et al., 2020, Wang et al., 2019). Herein, we hypothesize that the S2EBPR concept, where extended anaerobic phases to encourage hydrolysis and fermentation of sludge incorporated in the system, can be combined with short-cut N removal process to achieve simultaneous nitrogen and phosphorus removal for high strength wastewater. We propose that the combination of extended anaerobic phase and exposure to a mixture of relatively higher level of VFAs could potentially be an alternative strategy to achieve the nitrite-oxidizing bacteria (NOB) inhibition rather than the conventional strategies for suppressing NOB such as low DO manipulation, high free ammonia (FA) and free nitrous acid (FNA) inhibition and short SRT (Zhang et al., 2019).

In this study, we explore the feasibility of simultaneously coupled EBPR with short-cut N removal for treating synthetic high-strength wastewater mimicking the digester effluent of manure wastes. A lab-scale sequencing batch reactor (SBR) was applied and both the N and P removal performance were monitored by the routine chemical analysis of influent and effluent. Both phosphorus and nitrogen removal activities, including stoichiometry and kinetics, were evaluated via batch activity tests. The gene abundance of nitrifiers (AOB and NOB) were detected by real-time polymerase chain reaction (qPCR) and the microbial community were analyzed by 16S rRNA gene amplicon sequencing and oligotyping methods. To the best of our knowledge, this is the first study that reports the simultaneous high-efficient P removal and stable nitrite accumulation in one reactor for treating high-strength wastewater, which provides new insights and direction for effective and simultaneous N and P removal for treatment of centrate and other agricultural waste streams.

## 2. Materials and methods

### 2.1. SBR reactor operation

A lab-scale sequencing batch reactor (SBR) with a working volume of 4 L and a volumetric exchange ratio of 50% was used in this study (Fig. 1). The reactor was inoculated with EBPR activated sludge from Hampton Roads Sanitation District (HRSD) Virginia Initiative Treatment Plant (Norfolk, VA) and Nansemond Treatment Plant (Suffolk, VA). The reactor was operated with an 8 h cycle, consisting of a relatively long anaerobic phase (150 min, initial 6 min for feeding), 300 min aerobic phase, 15 min settling, 10 min decanting, and 5 min idle period resulting in a hydraulic retention time (HRT) of 16 h. During the whole process, pH is not controlled and varied between 7.2-8.5. Dissolved oxygen (DO) was maintained between 2-8 mg/L to avoid any oxygen limitations. The reactor was operated at a constant temperature of around 25 °C. The sludge retention time (SRT) was kept at 10 days by periodic wasting the mixed liquor at the end of the aerobic phase and MLVSS (mixed liquor volatile suspended solids) was maintained around 4500 mg/L.

**Fig 1.**
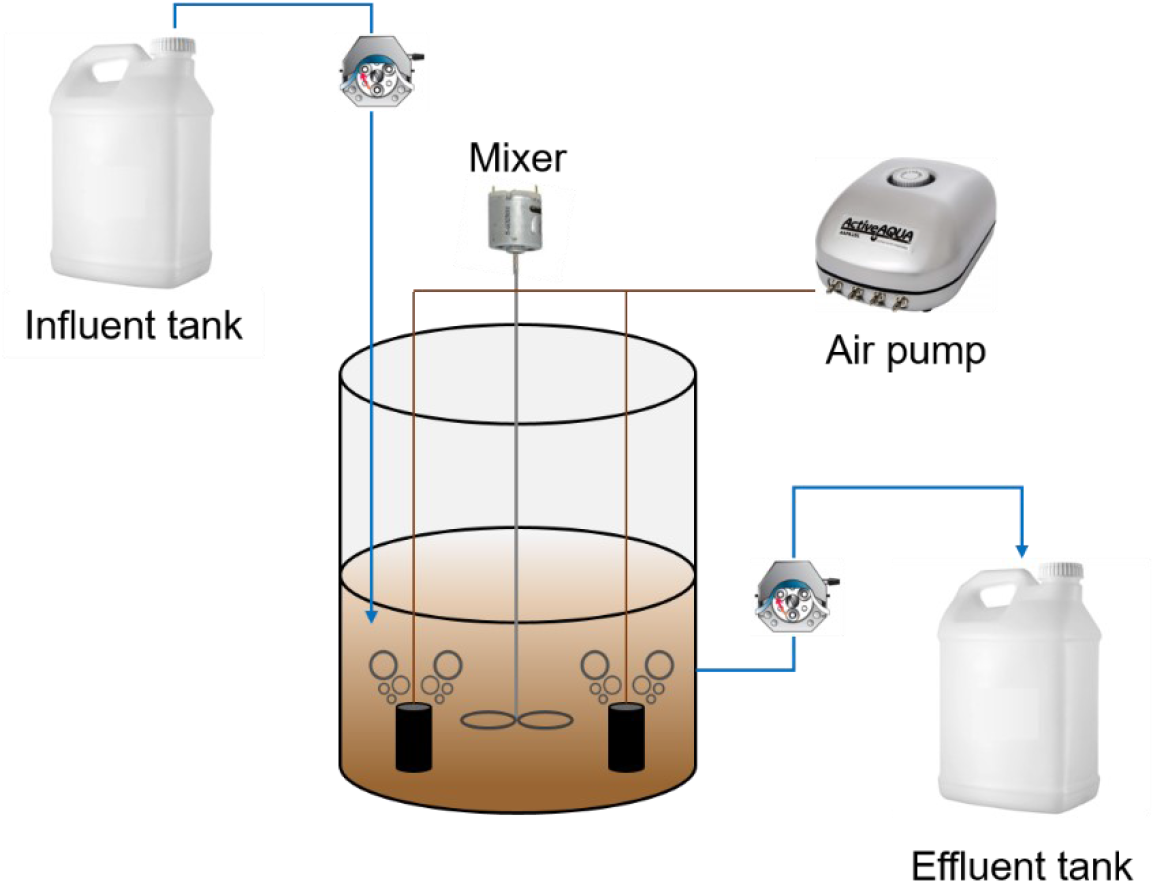
Configuration of SBR.

### 2.2. Synthetic wastewater

The synthetic wastewater was used in this study as the influent of SBR to simulate the digested manure wastewaters. The concentrations of COD, ammonium, and phosphate were determined according to the summary of the literature values (Table 1). The sodium acetate anhydrous (C_2_H_3_NaO_2_) and sodium propionate anhydrous (C_3_H_5_NaO_2_) were added as the carbon source with the molar ratio of 7:3. The combination of K_2_HPO_4_ and KH_2_PO_4_ (1.28:1 in molar ratio) was used as the phosphate source and NH_4_Q was the nitrogen source. The mineral medium of synthetic wastewater contained (per liter): 0.33 g MgSO_4_·7H_2_O, 0.15 g CaCl_2_·2H_2_O, 0.01 g ethylenediaminetetraacetic (EDTA), and 1.1 mL micronutrient solution (Marques et al., 2017). The micronutrient solution contained (per liter): 1.5 g FeCl_3_·6H_2_O, 0.15 g H_3_BO_3_, 0.03 g CuSO_4_·5H_2_O, 0.18 g KI, 0.12 g MnCl_2_·4H_2_O, 0.06 g Na_2_MoO_4_·2H_2_O, 0.12 g ZnSO_4_·7H_2_O, and 0.15 g CoCl_2_·6H_2_O (Smolders et al., 1994).

**Table 1.**
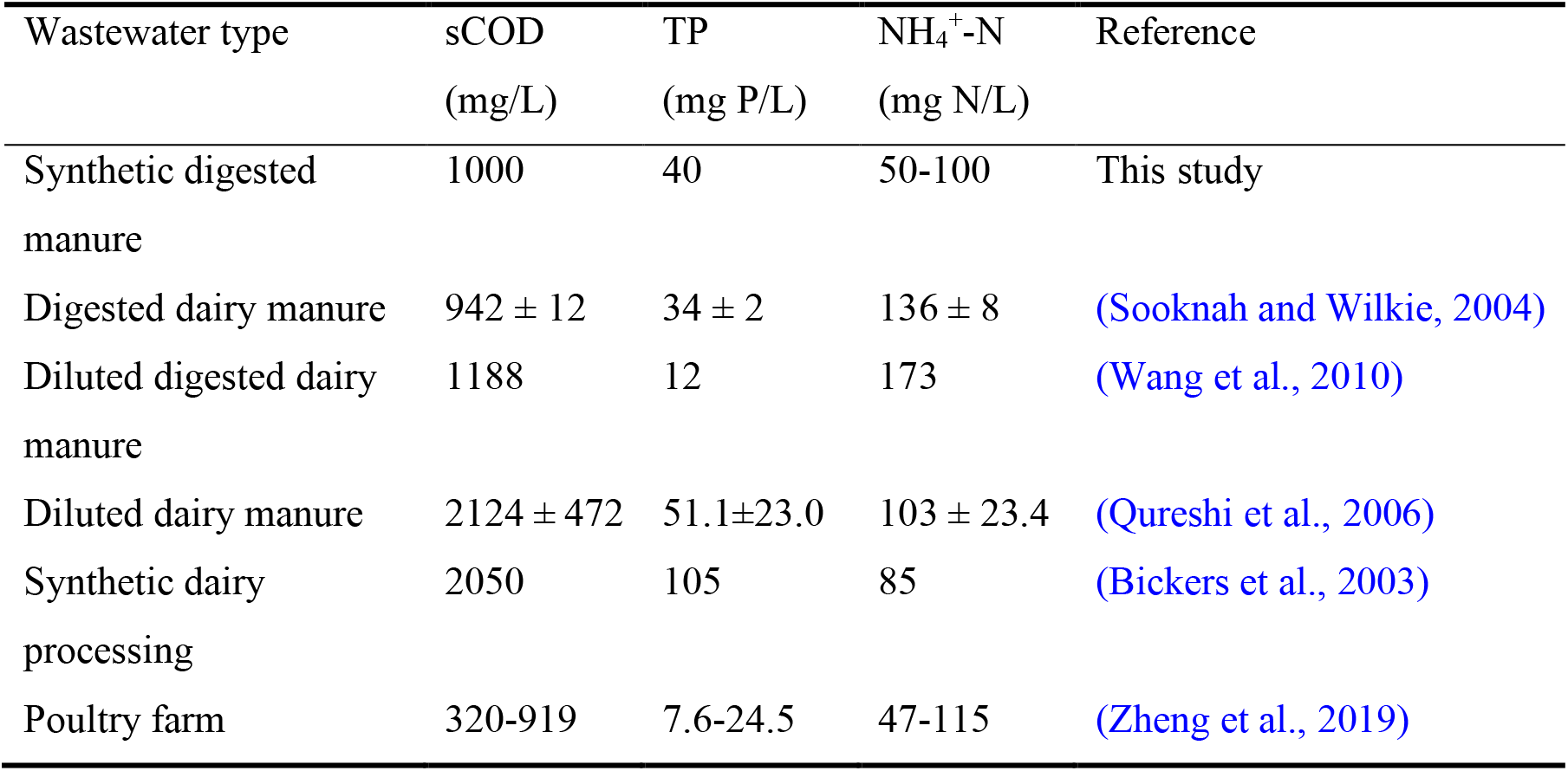
Summary of compositions and concentrations of manure wastewaters

### 2.3. Performance monitoring and chemical analysis

The influent and effluent of the reactor were sampled routinely and then filtered by 0.22 µm filters before chemical analysis. The concentrations of NH_4_^+^, NO_3_^-^, NO_2_^-^, and PO_4_^3-^ were measured by an ion chromatography (IC) (Thermo Fisher Scientific, USA) with Dionex ICS-2100 model. TOC and TN were measured by a total organic carbon analyzer (Shimadzu, Japan) with TOC-L model. COD, mixed liquor suspended solids (MLSS), and MLVSS were analyzed in accordance with Standard Methods (APHA, 2017). Nitrite accumulation ratio (NAR) was calculated as the ratio of NO_2_^-^-N to the sum of NO_2_^-^-N and NO_3_^-^-N (Regmi et al., 2014). Sludge samples were collected from the reactor at the end of the anaerobic and aerobic phases and centrifuged for 5 min with 6000 rpm. Poly-β-hydroxyalkanoates (PHAs) were extracted from freeze-dried sludge samples with a 3 h digestion time and 3% sulfuric acid and then analyzed by a gas chromatography-mass spectrometry (GC-MS) (Agilent, USA) with 7890A-5975C model (Lanham et al., 2013b). Glycogen was extracted from freeze-dried sludge samples with a 2 h digestion time and 0.9 M hydrochloric acid and then analyzed using the anthrone method for carbohydrate analysis (Zhang et al., 2015).

### 2.4. P release and uptake batch test

To evaluate the activity of PAOs in the sludge, the P release and uptake batch test was conducted *in situ* in the reactor for one cycle periodically following the previous procedures (Gu et al., 2008). During one entire cycle, water and sludge samples were collected at regular intervals to determine PHAs, glycogen, TOC, COD, TN, NH_4_^+^, NO_3_^-^, NO_2_^-^, PO_4_^3-^, MLSS, and MLVSS.

### 2.5. Nitrification batch activity test

To evaluate the nitrification activity of the sludge, batch activity tests were conducted according to previous methods (van Loosdrecht et al., 2016). Briefly, a certain amount of sludge (>1 g VSS/L) was collected at the end of the aerobic phase and washed with the mineral medium for three times. For the nitrification batch activity test, NH_4_Cl was added to achieve an initial ammonium concentration of 50 mg N/L, and the air was bubbled in to obtain the aerobic condition for 5 h. Samples were collected periodically throughout the entire tests to analyze TN, NH_4_^+^, NO_3_^-^, NO_2_^-^ and MLVSS. The reactor pH was continuously monitored and adjusted to maintain between 6.8 and 7.8 by adding 1 M NaOH or HCl. The temperature was maintained at 25 °C.

### 2.6. Nitrification batch inhibition test

To evaluate the impact of extended anaerobic exposure and high VFA concentration (as present in our SBR) on nitrification activity, batch tests were performed in duplicate. The sludge was taken from a lab-scale biological nutrients removal (BNR) reactor that performs full nitrification. Sludge was washed with the mineral medium first and split to conduct the following different batch tests. The control group included a 3 h aeration in order to obtain the normal nitrification rates of AOB and NOB. The experiment group included VFA (500 mg COD/L) addition at the beginning of a 2.5 h anaerobic starvation period, and then went through a 3 h aeration phase to assess the inhibition effects of the combination of extended anaerobic exposure and high VFA concentration. Samples were collected periodically throughout the entire tests to analyze TN, NH_4_^+^, NO_3_^-^, NO_2_^-^ and MLVSS.

### 2.7. DNA extraction and 16S rRNA gene amplicon sequencing

To investigate the microbial community dynamics, sludge samples were collected at the end of the aerobic phase periodically and the genomic DNA was extracted using the DNeasy PowerSoil Pro Kit (Qiagen, USA). The extracted DNA was sent to University of Connecticut-MARS facility for PCR amplification and sequencing targeting the V4 region using the primers 515F (5’-GTGCCAGCMGCCGCGGTAA-3’) and 806R (5’-GGACTACHVGG GTWTCTAAT-3’) and the amplicons were sequenced on the Illumina MiSeq with V2 chemistry using paired-end (2 x 250) sequencing. The raw paired-end reads were assembled for each sample and processed in Mothur v. 1.36.1 following the MiSeq standard operating procedure (SOP) (Kozich et al., 2013). High-quality reads were obtained after quality control and chimera screening and then clustered at a 97% similarity to obtain the operational taxonomic units (OTUs).

### 2.8. Quantitative polymerase chain reaction (qPCR)

The gene abundances of conventional autotrophic nitrifiers (AOB and NOB) were determined by qPCR using the CFX real-time PCR detection system (Bio-Rad, USA). The primer set of *amo*A gene of AOB was *amo*A-1F/2R (Rotthauwe et al., 1997). Thermocycling conditions were 95°C for 3 min, then 40 cycles of 95°C for 15 s, 54°C for 30 s, and 72°C for 60 s. The primer set of *nxrB* gene of *Nitrospira* was *nxr*B-169F/638R (Pester et al., 2014). Thermocycling conditions were 95°C for 5 min, then 35 cycles of 95°C for 40 s, 56.2°C for 40 s, and 72°C for 90 s. The qPCR reaction mixture in a total of 20 μL containing 1 μL DNA template, 1 μL each primer, 7 μL RNase-free water (Takara, Japan), and 10 μL iQ SYBR Green Supermix (Bio-Rad, USA). The standard curves were constructed from a series of 10-fold dilutions of plasmid DNA obtained by TOPO TA cloning (ThermoFisher, USA).

### 2.9. Oligotyping analysis and phylogenetic tree construction

Oligotyping was carried out to further resolve *Candidatus* Accmulibacter genus OTUs into sub-genotypes, namely oligotypes, based on the high-variational sites in reconstructed 16S sequences, following the protocol in previous studies (Eren et al., 2013, Srinivasan et al., 2019). To reveal the phylogeny of identified Accumulibacter oligotypes, a phylogenetic analysis was conducted on all identified oligotype representative sequences of our reactor (3 total), Accumulibacter oligotypes in a previous study from our group (9 total) (Srinivasan et al., 2019) and Accumulibacter phosphatis reference sequences in the MiDAS database (18 total) (McIlroy et al., 2015). An extra random sequence (*Dechloromonas*, FLASV96.1460) in the same *Rhodocyclaceae* family was chosen from the MiDAS database as outgroup. More details could refer to Supplementary Information (SI).

## 3. Results

### 3.1. Long-term performance of SBR

The SBR was running for 170 days in total and could be divided into three stages. In stage 1 (S1, day 1-105), the reactor started up with an average influent P concentration of 39.7 ±2.1 mg P/L, while the effluent P concentration was very unstable ranged from 3.5 to 23.1 mg P/L, with P removal efficiency fluctuated between 43.5%-88.2%. To evaluate the interference of nitrification on P removal, the average influent ammonium concentration was first set at 48.2 ± 6.5 mg N/L. Intriguingly, the nitrite accumulation was detected at around day 30. The effluent nitrite concentration could reach 6.0 ±2.4 mg N/L with no nitrate detected and the nitrite accumulation ratio (NAR) was nearly 100%. In stage 2 (S2, day 105-135), notably better and stable P removal performance was obtained with a higher average effluent P concentration of 2.4 ±1.3 mg P/L, resulting in P removal efficiency between 92.7%-99.2%. The effluent nitrite concentration was maintained at 7.3 ± 5.7 mg N/L, while nitrate was below the detection limit. In stage 3 (S3, day 135-170), the influent ammonium concentration was increased to around 100 mg N/L in reference to the digested manure wastewaters (Table 1). As a consequence, the effluent nitrite concentration increased to 20.4 ± 6.4 mg N/L with NAR still nearly 100%, while the average effluent P concentration was unaffected and further down to 0.8 ±1.0 mg P/L and the P removal efficiency was 99.5 ± 0.8%.

**Fig 2.**
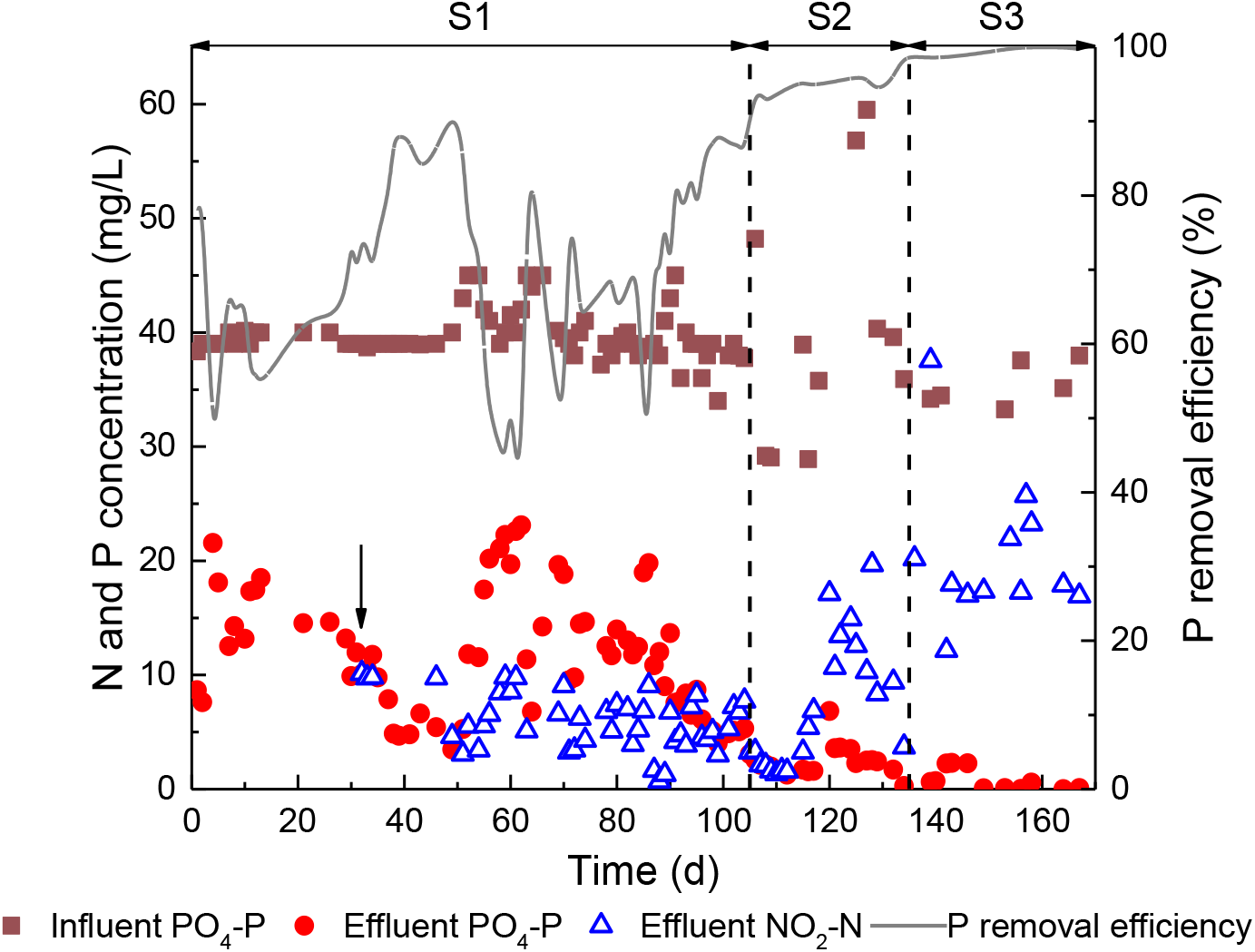
Long-term performance of SBR (the arrow indicated the time that effluent nitrite could be detected).

### 3.2. Phosphorus removal evaluation

#### 3.2.1. P release and uptake tests

To evaluate the EBPR metabolic activities in different stages, the *in-situ* variations of TOC, nutrients, PHAs, and glycogen in the typical SBR cycle were investigated. As shown in Fig. 3 and Fig. 4, both stage 2 (day 132) and stage 3 (day158) exhibited the typical PAO phenotypes of carbon uptake, P release, glycogen hydrolysis, PHA synthesis in the anaerobic phase, and P uptake, glycogen formation, PHA oxidation in the aerobic phase. During the first 30 min of the anaerobic phase, the input carbon source (195 ± 0.2 mg C/L of acetate and propionate) was almost taken up, and meanwhile, 133.5 mg P/L and 96.2 mg P/L was released in the typical cycle at S2 and S3, respectively. During the anaerobic phase for S2, PHA accumulated from 57.4 to 180.5 mg C/L, and 181.1 mg C/L glycogen was consumed to supply reducing power and energy. During the anaerobic phase for S3, a similar amount of glycogen was utilized and the PHA level increased from 77.3 to 118.1 mg C/L. In the subsequent aerobic phase for S2, an excess amount of phosphate was taken up resulting in 19.6 mg P/L net removal and the effluent P was 1.1 mg P/L. For S3, the net P removal declined to 10.9 mg P/L but the effluent P was only 0.6 mg P/L. Simultaneously, PHA was oxidized as the energy and carbon source for bacterial growth and glycogen replenishment, and the glycogen level increased from 600.1 mg C/L to 986.8 mg C/L for S2 and from 365.1 mg C/L to 727.2 mg C/L for S3. It was noted that ammonium was consumed in the first 90 min of the aerobic phase, leading to 8.7 mg N/L and 23.2 mg N/L of nitrite production for S2 and S3, respectively.

**Fig 3.**
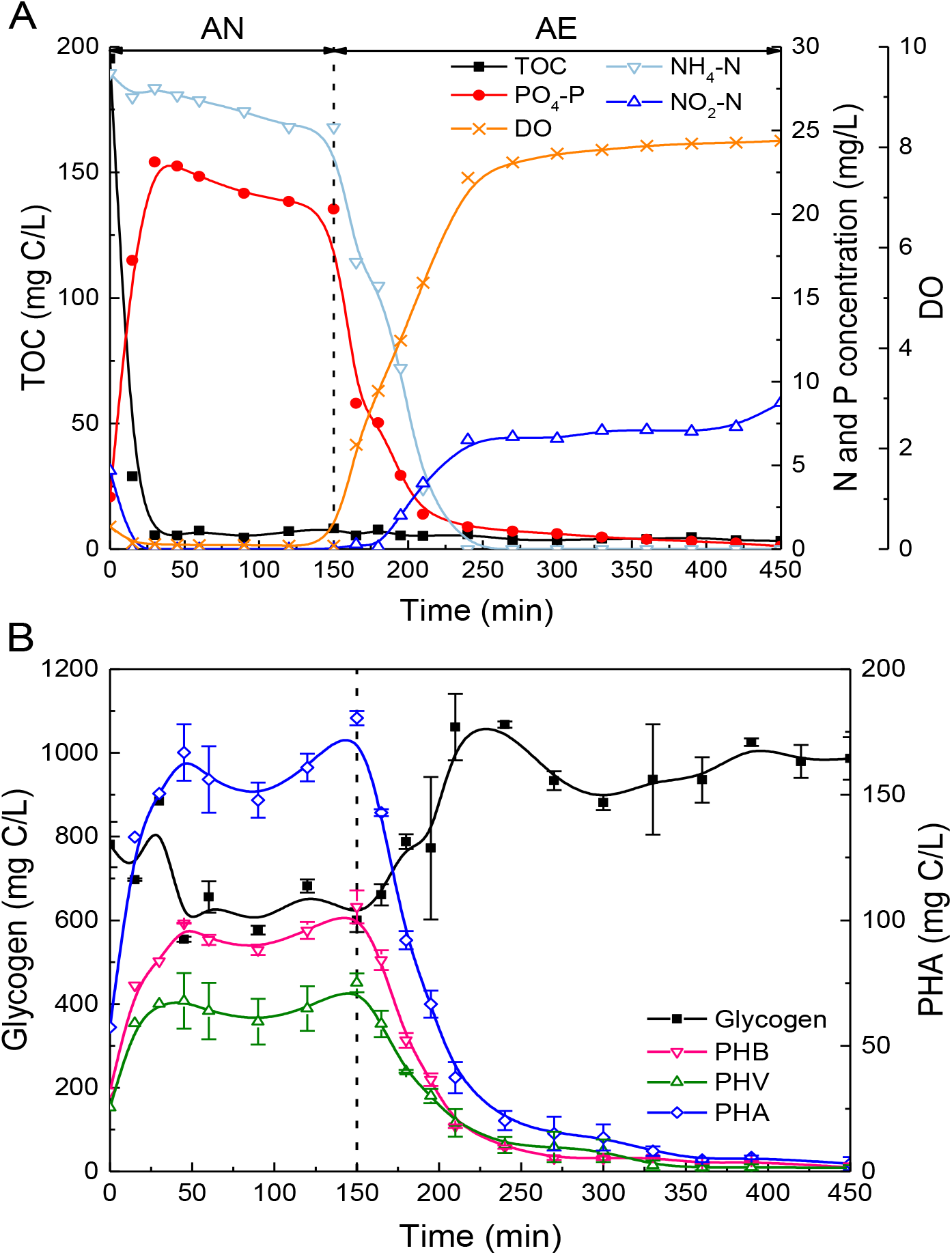
Profiles of (A) TOC, nitrogen, phosphorus, and DO concentrations; (B) glycogen and PHAs in typical cycle study at stage 2 (day 132). AN: anaerobic; AE: aerobic.

**Fig 4.**
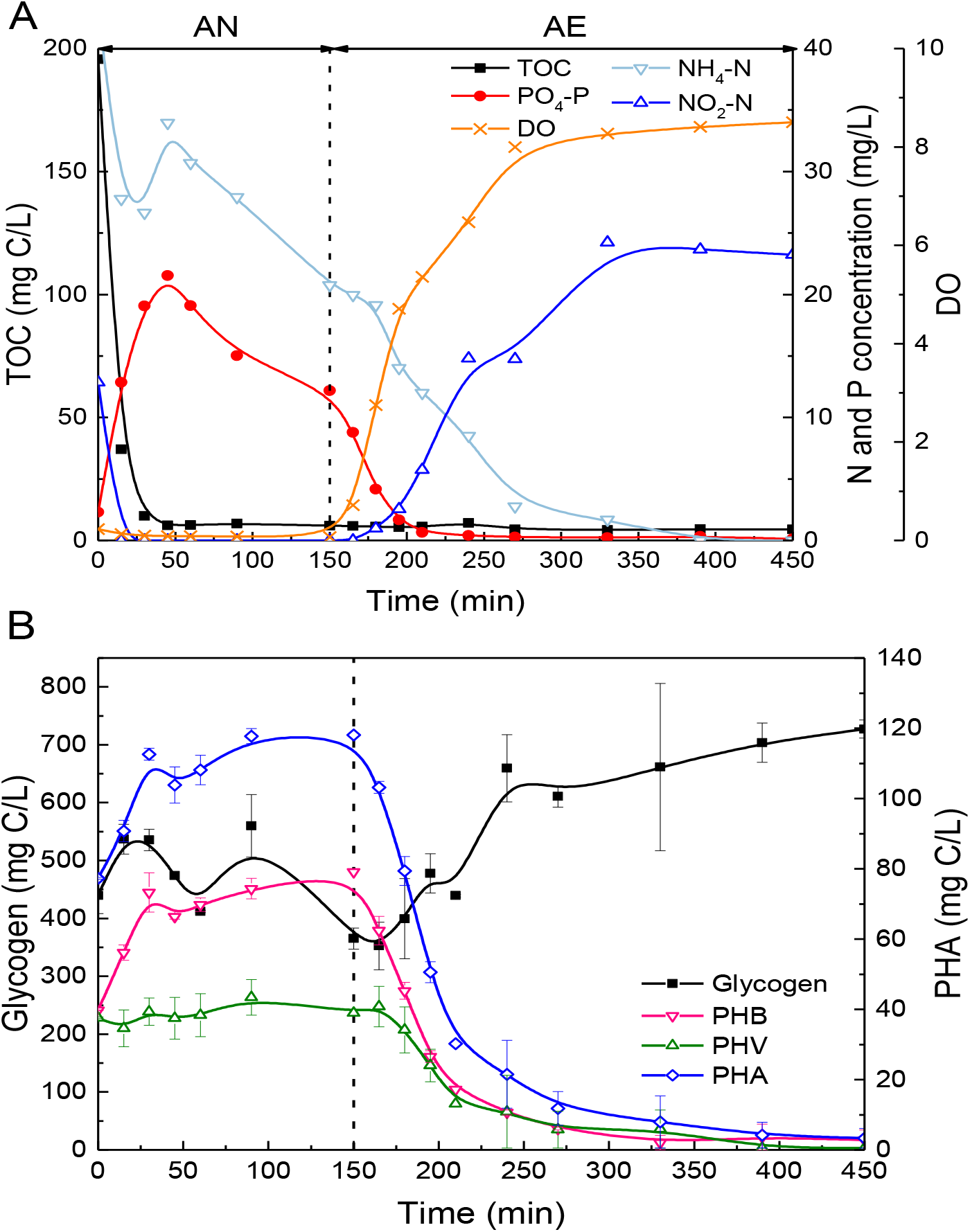
Profiles of (A) TOC, nitrogen, phosphorus, and DO concentrations; (B) glycogen and PHAs in typical cycle study at stage 3 (day 158). AN: anaerobic; AE: aerobic.

#### 3.2.2. EBPR kinetic and stoichiometric parameters

The kinetic and stoichiometry parameters were further computed to evaluate EBPR activity. The kinetic parameters of EBPR activities were compared with those reported in the literature (Table 2). The anaerobic VFA uptake rate, P release rate, and the aerobic P uptake rate were generally higher than those values observed in full-scale plants and were more comparable with the values reported in lab-scale systems. Especially, the anaerobic carbon uptake rate in our system fed with high levels of acetate and propionate was much higher than those in the conventional EBPR systems treating low-strength municipal wastewater (7.68-42.5 mg C/(g VSS·h)) but was at a similar magnitude with the values reported in a lab-scale acetate-fed EBPR system (233 mg C/(g VSS·h)). In stage 2, the P release and uptake rates were also comparable with the values obtained in the enriched PAOs culture (Oehmen et al., 2005a), indicating a high PAO activity and/or relative abundance, as observed in Section 3.2.3. In stage 3, however, the P uptake and release rates were halved, which might be related to inhibition associated with the significantly higher nitrite accumulation occurred in stage 3 than stage 2.

**Table 2.** The comparison of the kinetic rates in EBPR systems.

|  | VFA-Uptake<br>mg C/(g VSS·h)* | P-Release<br>mg P/(g VSS·h) | P-Uptake<br>mg P/(g VSS·h) | Reference |
| --- | --- | --- | --- | --- |
| Lab-scale (S2) | 85.14 | 60.77 | 27.64 | This study |
| Lab-scale (S3) | 85.35 | 31.04 | 13.94 | This study |
| Full-scale | 16.1-42.5 | 5.6-31.9 | 2.4-9.7 | (Gu et al., 2008) |
| Full-scale | - | 11.1-19.2 | 1.9-11 | (He et al., 2008) |
| Full-scale | 7.68-24.9 | 2.85-5.3 | 0.6-2.6 | (Onnis-Hayden et al., 2020) |
| Lab-scale | 233 | 88.8 | 27.3 | (Oehmen et al., 2005a) |
| Lab-scale | 7.71-32.66 | 4.38-50.63 | 9.82-23.81 | (Majed and Gu, 2020) |

\* The VFA uptake has been corrected excluding denitrification consumption.

The stoichiometric ratios were also calculated to evaluate the EBPR activity (Table 3). The carbon utilization due to nitrite denitrification was deducted for EBPR activities evaluation. The P release to VFA uptake ratio (P/VFA) is regarded as the indicator of the relative abundance and activities of PAOs and GAOs and carbon utilization efficiency (Schuler and Jenkins, 2003). The P/VFA ratio in stage 2 and stage 3 were 0.27 P-mol/C-mol and 0.21 P-mol/C-mol, respectively. These values were between the value of PAO and GAO model (Table 3), indicating the presence of GAOs in the system, which was consistent with microbial community results (Section 3.2.3). The aerobic P uptake to PHA utilization ratio (P/PHA) in both stage 2 and stage 3 was around 0.33-0.36 P-mol/C-mol that was close to the predicted value for the PAO model with enriched *Candidatus* Accumulibacter, which was consistent with the observed dominance of *Candidatus* Accumulibacter in our reactor. The P/VFA and P/PHA ratios together indicated the coexistence and co-prosperity of both PAOs and GAOs, which were further confirmed by 16S rRNA gene amplicon sequencing (Section 3.2.3).

**Table 3.**
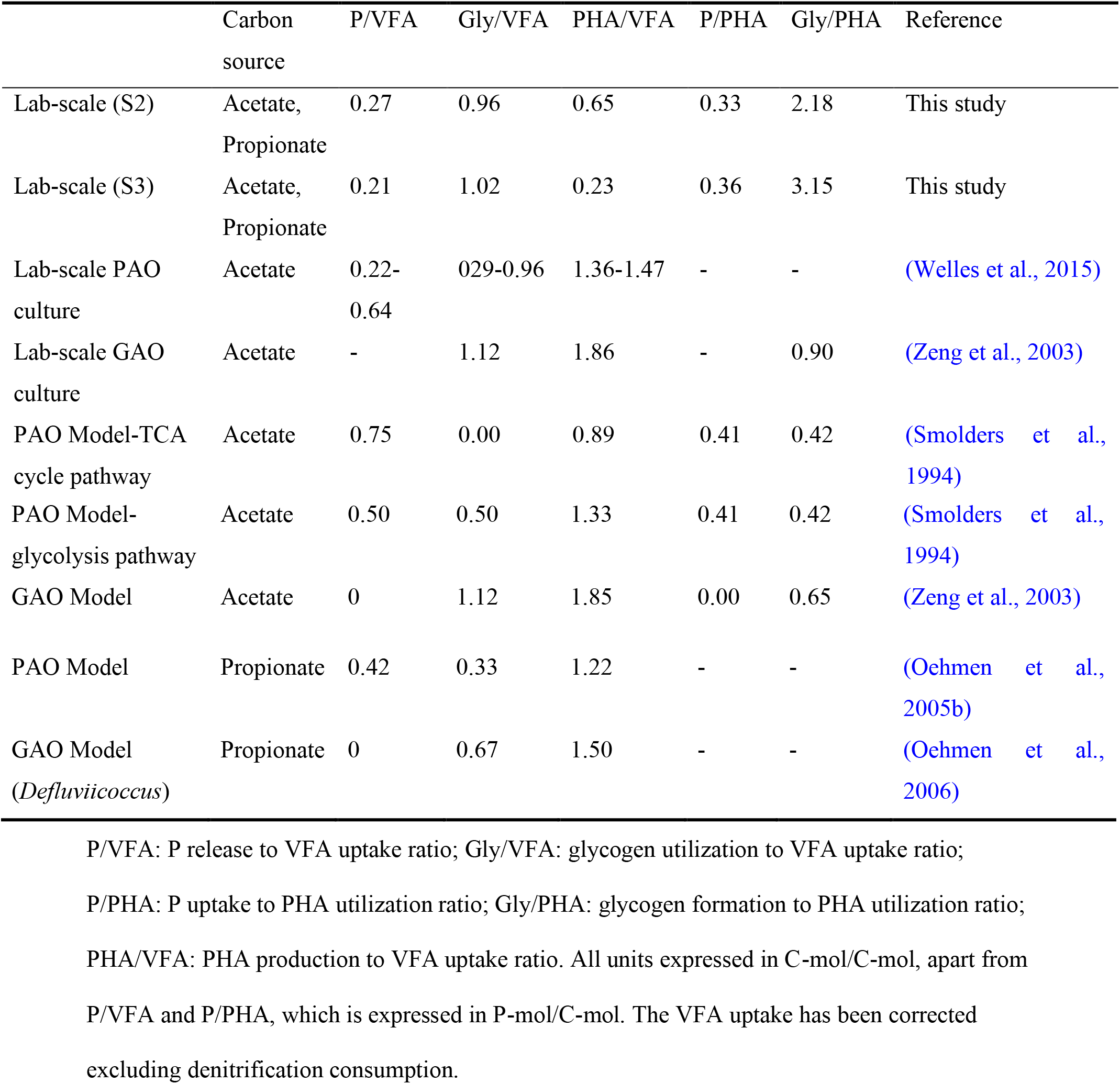
The comparison of the stoichiometric ratios of EBPR system.

|  | Carbon source | P/VFA | Gly/VFA | PHA/VFA | P/PHA | Gly/PHA | Reference |
| --- | --- | --- | --- | --- | --- | --- | --- |
| Lab-scale (S2) | Acetate, Propionate | 0.27 | 0.96 | 0.65 | 0.33 | 2.18 | This study |
| Lab-scale (S3) | Acetate, Propionate | 0.21 | 1.02 | 0.23 | 0.36 | 3.15 | This study |
| Lab-scale PAO culture | Acetate | 0.22-0.64 | 0.29-0.96 | 1.36-1.47 | - | - | (Welles et al., 2015) |
| Lab-scale GAO culture | Acetate | - | 1.12 | 1.86 | - | 0.90 | (Zeng et al., 2003) |
| PAO Model-TCA cycle pathway | Acetate | 0.75 | 0.00 | 0.89 | 0.41 | 0.42 | (Smolders et al., 1994) |
| PAO Model-glycolysis pathway | Acetate | 0.50 | 0.50 | 1.33 | 0.41 | 0.42 | (Smolders et al., 1994) |
| GAO Model | Acetate | 0 | 1.12 | 1.85 | 0.00 | 0.65 | (Zeng et al., 2003) |
| PAO Model | Propionate | 0.42 | 0.33 | 1.22 | - | - | (Oehmen et al., 2005b) |
| GAO Model<br>(*Defluviicoccus*) | Propionate | 0 | 0.67 | 1.50 | - | - | (Oehmen et al., 2006) |

P/VFA: P release to VFA uptake ratio; Gly/VFA: glycogen utilization to VFA uptake ratio; P/PHA: P uptake to PHA utilization ratio; Gly/PHA: glycogen formation to PHA utilization ratio; PHA/VFA: PHA production to VFA uptake ratio. All units expressed in C-mol/C-mol, apart from P/VFA and P/PHA, which is expressed in P-mol/C-mol. The VFA uptake has been corrected excluding denitrification consumption.

The anaerobic glycogen utilization to acetate uptake ratio (Gly/VFA) can provide insights into the extent of utilization of glycolysis compared to the tricarboxylic acid (TCA) cycle for typical PAO/GAO metabolism in the EBPR system. Typically, known GAOs solely rely on glycogen as the energy source via the glycolysis pathway for anaerobic PHA synthesis, while known PAOs can utilize polyP as an alternative. Therefore, the theoretical Gly/VFA ratio for known GAOs is higher (typically over 0.7) than those predicted for PAO (Smolders et al., 1994). In this study, Gly/VFA ratio in both stages was over 0.95 and close to the GAO model, indicating the dominant involvement of GAO metabolism. Meanwhile, considering the high abundance of known PAOs in the system revealed by sequencing, this high Gly/VFA ratio might also be associated with the preferential use of glycolysis pathway over TCA cycle for PAOs, which was observed in side-stream hydrolysis and is considered more efficient through producing additional PHA and releasing less phosphate per substrate uptake, thus potentially beneficial for EBPR (Lanham et al., 2013a, Wang et al., 2019).

The anaerobic PHA generation to VFA uptake ratios (PHA/VFA) in stage 2 and stage 3 was 0.65 C-mol/C-mol and 0.23 C-mol/C-mol, respectively, both of which were much lower than those theoretical values predicted for either PAO or GAO models (1.33 and 1.85 respectively). This indicated that there were possibly other carbon uptake and storage mechanisms beyond the current understanding, warranting further investigation. A possible explanation for that could be due to the sufficient carbon input in our system that promoting not only known PAOs and GAOs, but also unknown ones as well as the other heterotrophic microorganisms with intracellular carbon, such as PHA-producing capabilities as evidenced by our previous study (Wang et al., 2018). It was also noticed that the aerobic glycogen formation over PHA consumption ratios (Gly/PHA) was 2.18 and 3.15 for S2 and S3, respectively, which were significantly higher than any theoretical model predictions (0.42 to 0.90). These further suggest that there were possibly other intracellular carbon storage compounds and/or associated unrecognized pathway for glycogen formation.

#### 3.2.3. Microbial community of PAOs

The microbial communities were characterized by 16S rRNA gene amplicon sequencing (Fig. 5). For known candidate PAOs and GAOs in the SBR during different stages according to the MiDAS database (McIlroy et al., 2015), the canonical *Candidatus* Accumulibacter was the dominant genus of known PAOs, and its relative abundance increased from 0.6% (day 32) to 33.1% (day 144), which was consistent with the improved and steady EBPR performance. Another candidate PAO, namely *Tetrasphaera*, was below the detection limit at all stages in the system (Nielsen et al., 2019). The relative abundance of *Acinetobacter* was 1.2%, which has been reported in the SBR reactor as the potential PAO (Seviour et al., 2003, Nguyen et al., 2011). *Dechloromonas* was proved to be a potential PAO (Petriglieri et al., 2019), but its relative abundance was below 1%. Furthermore, oligotyping analysis was performed to reveal the microdiversity within Accumulibacter OTUs and 3 oligotypes of Accumulibacter were identified. The relative abundance of oligotype 2 was up to 88.2% at the startup of the reactor (Figure 6A). Oligotype 1 became the dominant sub-type, accounting for 64.2-87.9% of the total Accumulibacter, (Fig. 6A). By phylogenetic comparison with the previous study from our group and the literature, the results revealed that the dominant oligotype 1 in our reactor was highly consistent with the specific Accumulibacter oligotype 2 (belonging to Clade IIC) accounting for 62.4-68.0% in the full-scale side-stream EBPR system (highlighted in Fig. 6B), which was observed to be much lower (<20%) in the conventional EBPR-A2O system (Srinivasan et al., 2019).

**Fig 5.**
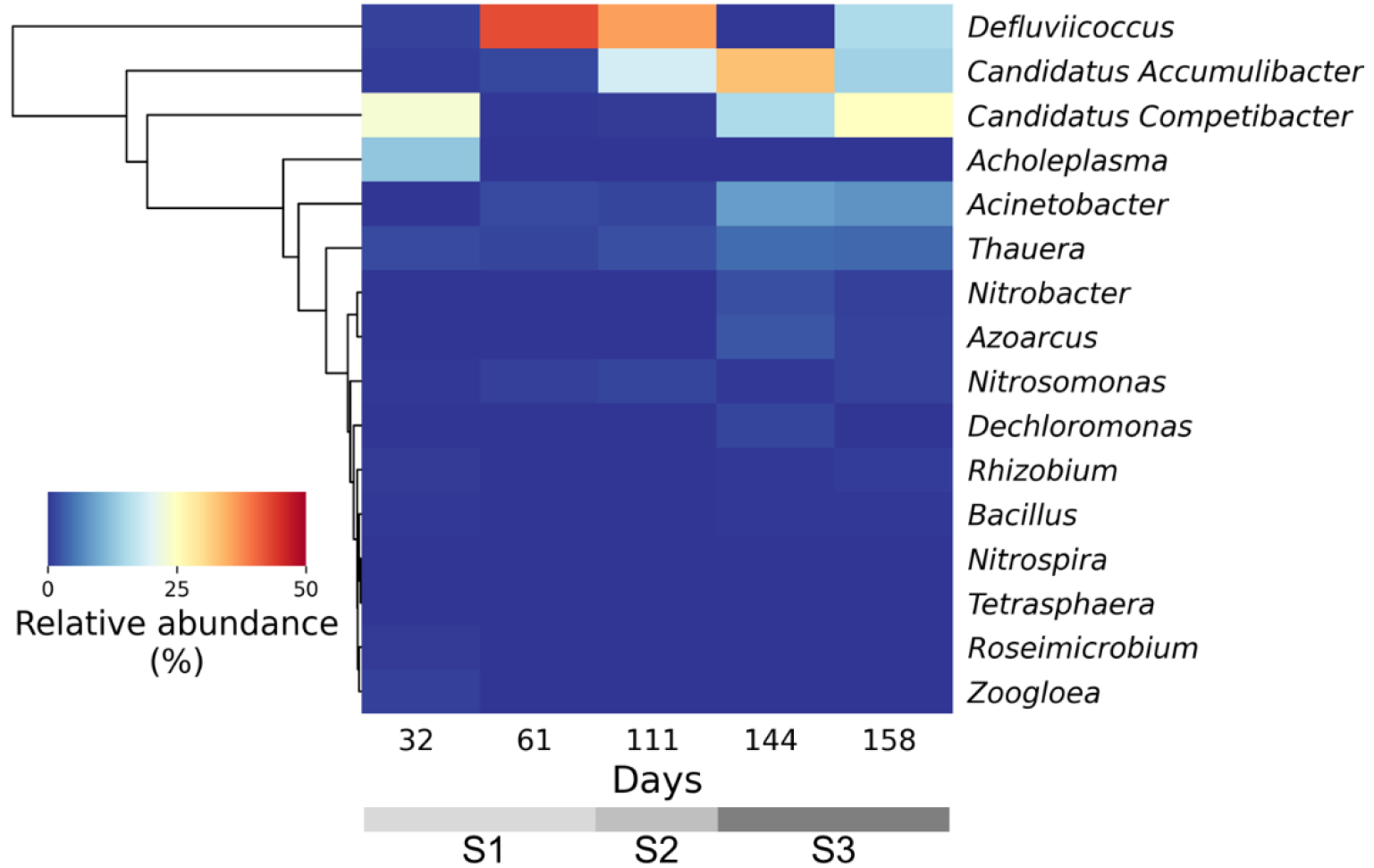
Heatmap of microbial community at genus level at different stages of the lab-scale reactor operation.

**Fig 6.**
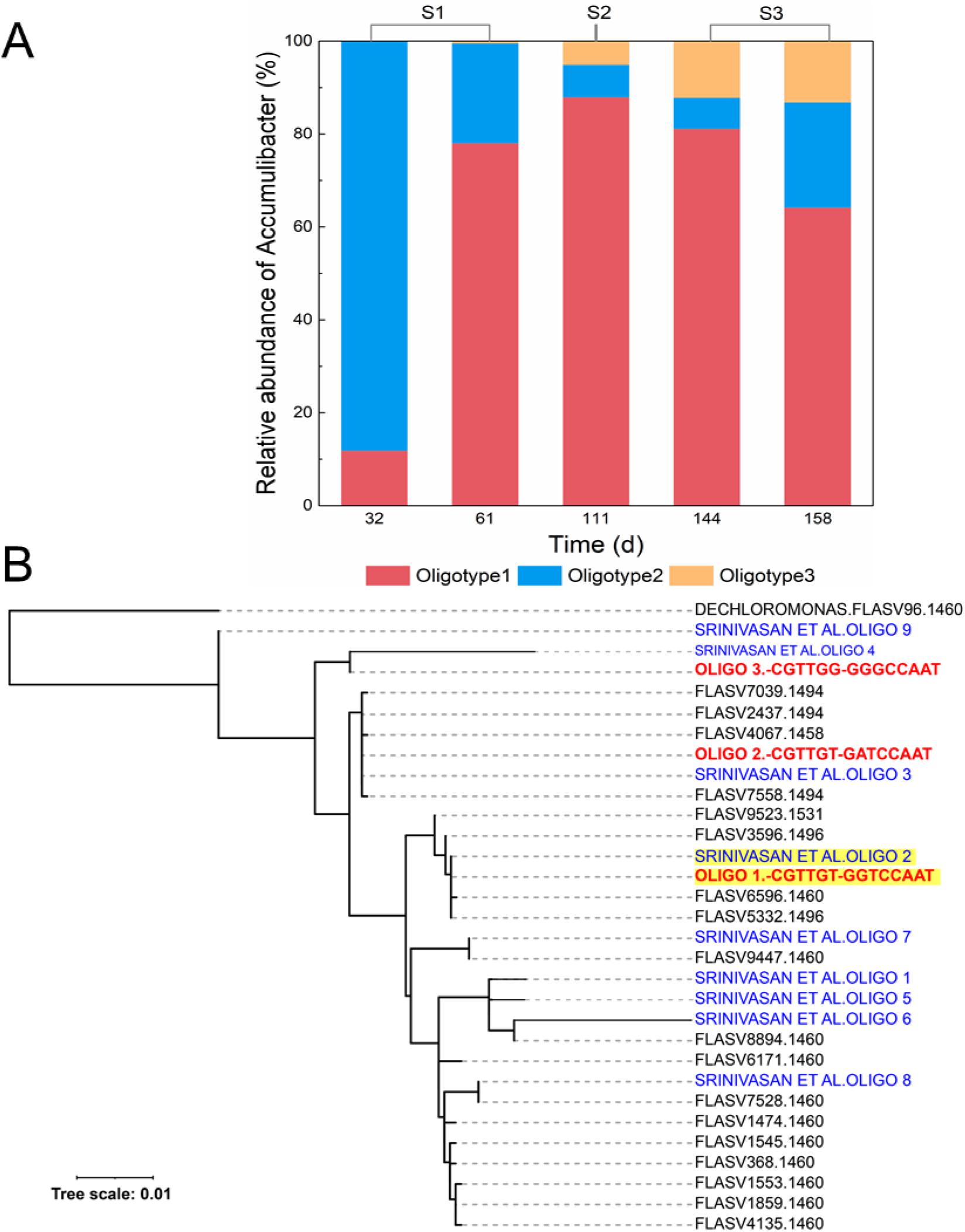
(A) Relative abundance of oligotypes of 16S rRNA gene amplicon sequences classified as *Candidatus* Accumulibacter at different stages of the lab-scale reactor operation; (B) Phylogenetic tree of *Candidatus* Accumulibacter oligotype representative sequences in our system (in red), oligotypes curated in the previous study from full-scale side-stream EBPR sludge (Srinivasan et al., 2019) (in blue) and reference sequences of *Candidatus* Accumulibacter phosphatis from the MiDAS database (McIlroy et al., 2015) (in black).

The GAOs are known to compete with PAOs that can impact the EBPR performance (Schuler and Jenkins, 2003, Gu et al., 2008). *Defluviicoccus* was found to be the most dominant GAO in the system with a relative abundance of 36.0-42.6% at stage 2 (Wong et al., 2004). The predominance of *Defluviicoccus* was proved to be highly associated with the higher relative ratio of propionate over acetate in the feed because its propionate uptake rate is comparable to *Candidatus* Accumulibacter and much higher than *Candidatus* Competibacter (Dai et al., 2007). Compared to stage 2, the known GAO populations shifted to *Candidatus* Competibacter over *Defluviicoccus* with a relative abundance of 15.3-24.9% at stage 3. Factors that govern the changes in known GAO community composition remain unclear including SRT (Onnis-Hayden et al., 2020), C/P ratios (Majed and Gu, 2020) and carbon compositions (Shen and Zhou, 2016) have been evidenced to be associated with GAOs shifts. The accumulation of *Defluviicoccus* in the system may be due to the propionate in the feeding medium, which prefers propionate instead of acetate in order to maximize their production of PHAs with the same glycogen consumption (Dai et al., 2007).

### 3.3. Nitrogen removal evaluation

#### 3.3.1. Nitrification batch activity test

The nitrification batch activity tests were performed with the sludge taken from the different stages to determine the maximum activity of AOB and NOB in the SBR. The results showed that the ammonia oxidizing rate (AOR) and nitrite oxidation rate (NOR) were as low as 8.46±1.14 mg N/(g VSS·d) and 0.35±0.09 mg N/(g VSS·d), respectively in stage 2 (Fig. S3). While with the increase of the influent ammonium concentration in stage 3, both the AOR and NOR were enhanced to 27.42±4.25 mg N/(g VSS·d) and 4.73±1.61 mg N/(g VSS·d) (Fig. S4), but still was much lower to the normal value reported in the literature (Table 4). According to the nitrogen mass balance of *in situ* batch activity tests (Fig. 3), around 8 mg N/L was removed in the anaerobic phase via denitrification, 18.1 mg NH_4_-N/L (67.5% of total ammonium) was estimated for biomass synthesis and the rest of 8.7 mg NH_4_-N/L (32.5% of total ammonium) was oxidized to nitrite in the aerobic phase (See calculations in SI). The nitrification batch activity test indicated that the partial nitrification was achieved in the system with the inhibition of NOB activity.

**Table 4.** The comparison of the nitrification rates of the activated sludge system.

| Scale | AOR [mg N/(g VSS·d)] | NOR [mg N/(g VSS·d)] | Reference |
| --- | --- | --- | --- |
| Lab-scale (S2) | $8.46 \pm 1.14$ | $0.35 \pm 0.09$ | This study |
| Lab-scale (S3) | $27.42 \pm 4.25$ | $4.73 \pm 1.61$ | This study |
| Lab-scale | 86.88 | 133.68 | (Li et al., 2018) |
| Full-scale | 85.4 | 138 | (Seuntjens et al., 2018) |
| Full-scale | 78 | 112.25 | (Yao and Peng, 2017) |

AOR: ammonia oxidizing rate; NOR: nitrite oxidizing rate.

#### 3.3.2. Nitrification batch inhibition test

The comparatively lower AOB and NOB activities observed raised question on whether the extended anaerobic exposure and high VFA concentration could exert inhibition on both AOB and NOB and if it can lead to or contribute to the NOB out-selection. To further investigate and probe the reasons for NOB out-selection in the SBR, a series of nitrification inhibition batch tests, using sludge from a lab-scale biological nutrients removal (BNR) reactor that performs efficient nitrification, were performed to evaluate the impact of anaerobic starvation and high VFA concentration on nitrification rates. The results showed that the control AOR and NOR were 164.97 ± 0.54 mg N/(g VSS·d) and 84.21 ± 1.27 mg N/(g VSS·d), respectively. However, with the combination of extended anaerobic exposure and high concentration of VFAs similar to those in our reactor, there was only a 3.7% decrease of AOR but over 10% decrease of NOR. The results suggested that NOB were more severely inhibited by VFA than AOB (Table 5), which most likely contributed to the NOB suppression and consequent nitrite cumulation for over 140 days (over 420 cycles) in our system.

**Table 5.**
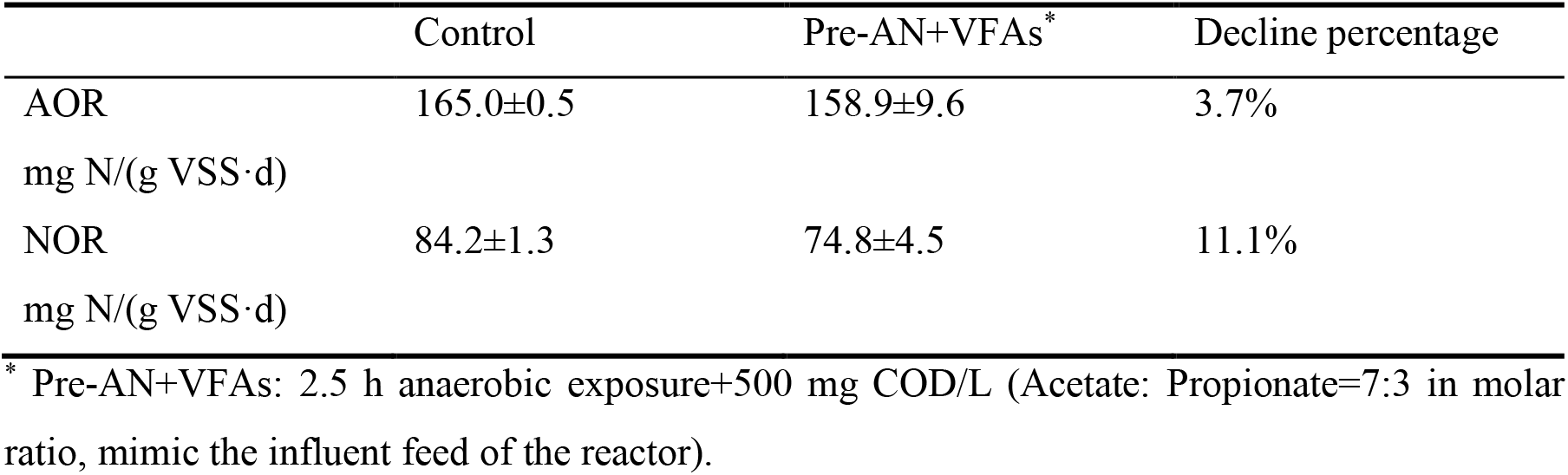
Nitrification rate under different treatment conditions.

#### 3.3.3. Microbial community of nitrifiers

The gene abundance of AOB and NOB was detected by both 16S rRNA gene sequencing and qPCR to further prove the NOB out-selection for all three stages. Based on the 16s rRNA gene sequencing analysis (Fig. 5), *Nitrosomonas* was the only known AOB genus that could be detected with the relative abundance of only (0.8±0.4)%, which was consistent with the low nitrification rate (Table 4) and NOB relative abundance was below the detection limit (<0.1%). qPCR results proved that the average *amo* gene abundance was [(6.98±2.12)×10^9^ copies/g] in stage 1 and stage 2, which was one order higher than *nxr* gene abundance [(2.18±1.50)×10^8^ copies/g] (Fig. 7). The average ratio of *amo*/16S was (1.7±0.7)% and *nxr*/16S ratio was less than 0.1%. In stage 3, both the gene abundance decreased, but *amo* gene abundance [(3.68±2.10)×10^8^ copies/g] was still maintained one order higher than *nxr* gene abundance [(7.56±0.87)×10^7^ copies/g], which were consistent with nitrification rates and the nitrite accumulation effect. The PCR results of key biomarkers indictive of AOB and NOB activities are consistent with the nitrification rates evaluation and system performance.

**Fig 7.**
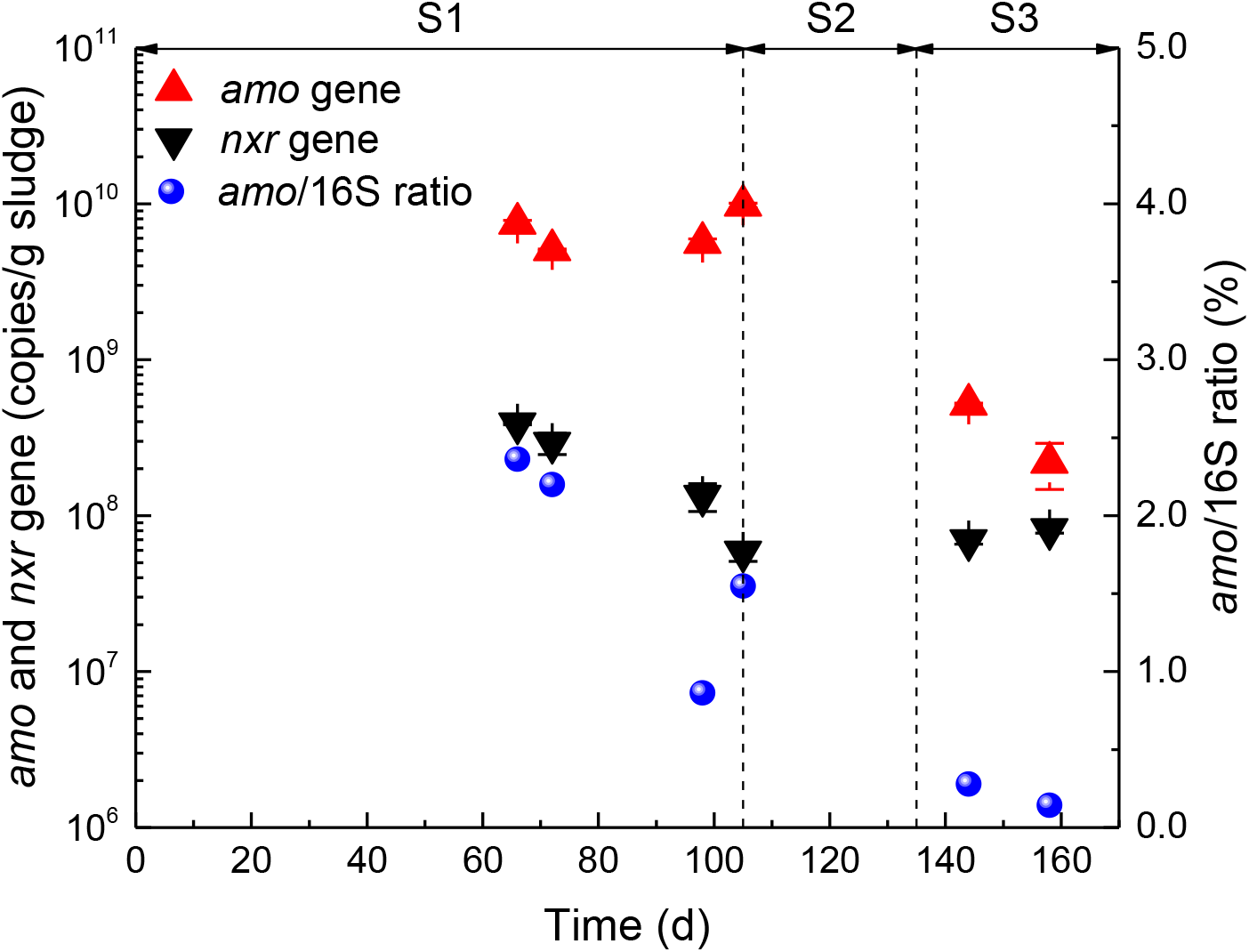
Relative abundance of AOB and NOB in SBR at different stages of the lab-scale reactor operation.

## 4. Discussion

### 4.1. New insights in the application of EBPR system for high-strength wastewater

The EBPR process, as one of the economic and high-efficient biological phosphorus removal techniques, has been widely applied for the treatment of domestic wastewater with the typical P concentrations between 4 and 12 mg P/L (Metcalf et al., 1979). However, the knowledge of applying EBPR in high-strength industrial and agricultural wastewaters (9-280 mg P/L) is still very limited (Ashekuzzaman et al., 2019). Therefore, in this study, the high-strength P concentration of 40 mg P/L with the rbCOD: P ratio as 25:1 was applied to test the feasibility and efficacy of the EBPR applications for high-strength wastewater such as agricultural manure digestate. Long-term reactor operation showed that the excellent P removal performance could be achieved in stage 2 and stage 3 of the lab-scale SBR. Especially in stage 3 (35 days with 105 cycles), the effluent P was as low as 0.8 ± 1.0 mg P/L and the P removal efficiency was up to 99.5 ± 0.8%, which is superior to the previous studies treating the high-strength manure wastewater (Table 6).

**Table 6.**
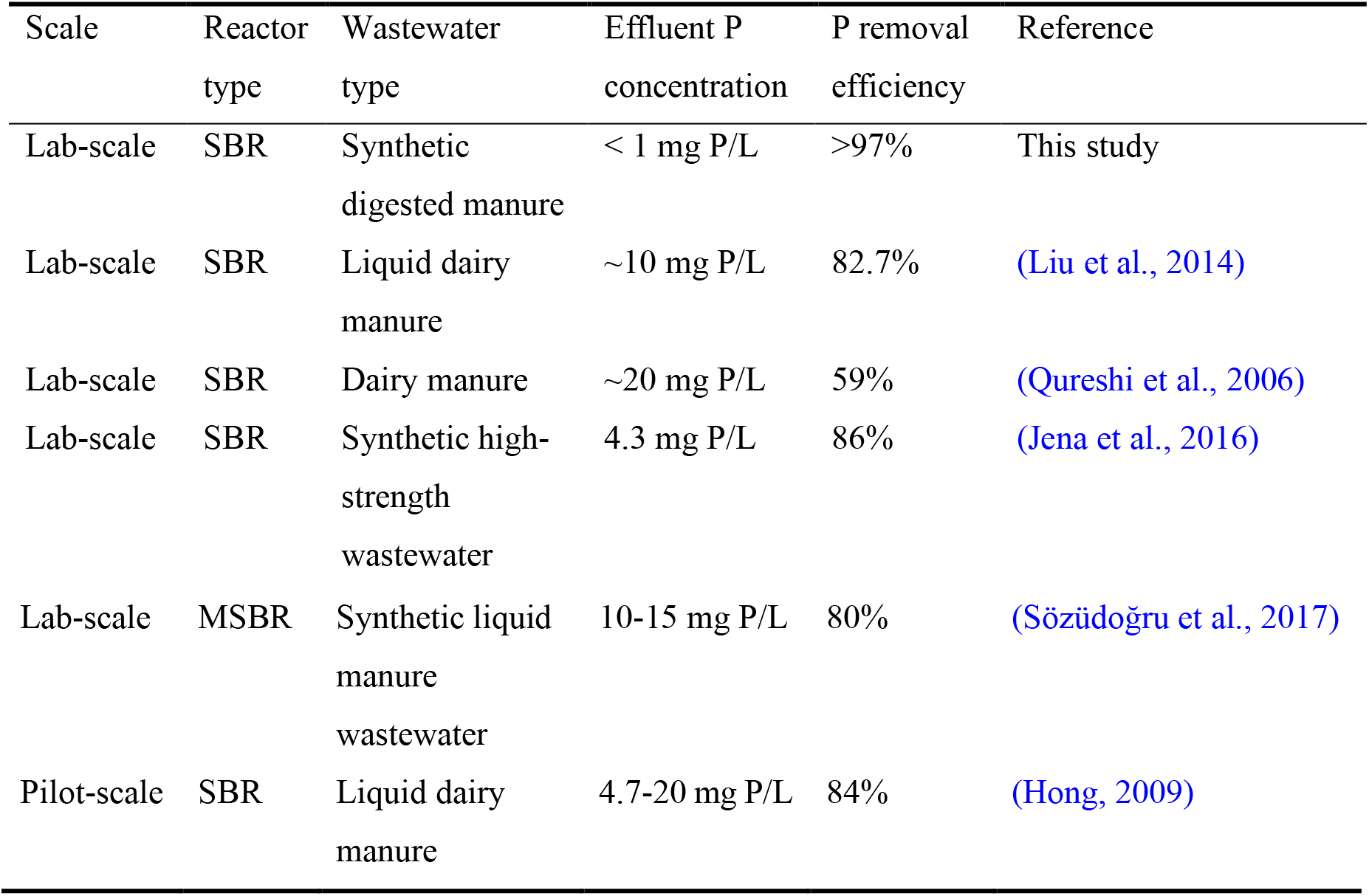
The comparison of the P removal performance of high-strength wastewater.

| Scale | Reactor type | Wastewater type | Effluent P concentration | P removal efficiency | Reference |
| --- | --- | --- | --- | --- | --- |
| Lab-scale | SBR | Synthetic digested manure | < 1 mg P/L | >97% | This study |
| Lab-scale | SBR | Liquid dairy manure | ~10 mg P/L | 82.7% | (Liu et al., 2014) |
| Lab-scale | SBR | Dairy manure | ~20 mg P/L | 59% | (Qureshi et al., 2006) |
| Lab-scale | SBR | Synthetic high-strength wastewater | 4.3 mg P/L | 86% | (Jena et al., 2016) |
| Lab-scale | MSBR | Synthetic liquid<br>manure<br>wastewater | 10-15 mg P/L | 80% | (Sözüdoğru et al., 2017) |
| Pilot-scale | SBR | Liquid dairy<br>manure | 4.7-20 mg P/L | 84% | (Hong, 2009) |

Interestingly, during the operation, nitrite gradually accumulated and the effluent nitrite could reach up to 20.4 ± 6.4 mg N/L. Numerous studies have reported that the presence of nitrite in the range of 10-20 mg/L would cause 3.0-17.6% inhibition on aerobic P uptake rate for different sludge type (Jabari et al., 2016). The specific batch test proved that 1 mg N/L nitrite could decrease the 80% aerobic P uptake activity and the activity was almost completely lost at 12 mg N/L (Saito et al., 2004). Series of batch experiments were performed to investigate the effect of nitrite on a highly enriched culture of *Candidatus* “Accummulibacter phosphatis” and 50% inhibition occurred at 2.0 mg NO_2_^-^-N/L and pH 7.0 (Pijuan et al., 2010). However, in this study, the presence of nitrite didn’t deteriorate the EBPR performance, even with the increase of nitrite from S2 (7.3 ±5.7 mg N/L) to S3 (20.4 ± 6.4 mg N/L), the P removal efficiency was enhanced from 95.1 ± 1.8% to 99.5 ± 0.8%. Moreover, the relative abundance of *Candidatus* Accumulibacter in this system maintained a stable level around 14.2-33.1% at S2 and S3. This suggest that certain PAOs or Accumulibacter sub-types can be adapted to higher nitrite level than previously assumed, and it is possible to design a system that involves both EBPR and nitrite accumulation for short-cut N removal. As discussed in section 3.2.3, the Accumulibacter oligotype in this study was highly phylogenetically consistent with the dominant one enriched in side-stream EBPR system, where the working condition was similar to our system, which provide different selective pressure and conditions that are quite different from those in the conventional activated sludge and EBPR systems.

Except for the nitrite inhibition, another key factor affecting EBPR performance is the PAO/GAO competition. The high readily biodegradable COD concentration in the influent with relatively high COD/P ratio would imply a higher relative abundance of GAOs (Schuler and Jenkins, 2003, Gu et al., 2008). As expected, both EBPR activities assessment and the microbial population analysis showed that a very stable and good EBPR performance was achieved in the presence of a relatively higher abundance of GAOs (*Defluviicoccus* and *Candidatus* Competibacter) (15.6-40.3%) than PAOs (*Candidatus* Accumulibacter) (14.2-33.1%). This was consistent with the previous understanding that the presence GAOs does not necessarily lead to EBPR deterioration, as long as the condition is kinetically favoring PAOs over GAOs (Gu et al., 2008, Nielsen et al., 2019). Although most of the stoichiometric parameters during the anaerobic phase of the SBR suggested dominant traits of GAO models, P uptake to PHA utilization ratio in aerobic phase (P/PHA) which can represent the ability of utilization of PHA for PAOs (0.33-0.36) was very close to the PAO model (0.41), indicating effective EBPR activities.

### 4.2. Potential new strategy for NOB out selection in systems that implement EBPR

In this study, both nitrification activity tests and microbial population analysis indicated that NOB was successfully suppressed and out-selected and stable nitrite accumulation was achieved in the SBR with simultaneous EBPR for 140 days (420 cycles). So far, several common factors have been employed to facilitate NOB out-selection to achieve partial nitrification including pH, DO, temperature, free ammonia (FA), and free nitrous acid (FNA) (Park et al., 2010, Zhang et al., 2019). Those control strategies are mostly successfully applied in high-strength ammonium-containing wastewater, but for mainstream conditions, the stable and effective suppression of NOB is still challenging (Cao et al., 2017). In our system, the DO in the aerobic phase was not controlled and close to the saturation level (>5 mg/L) (Fig. 4), while low DO below 1 mg/L is usually controlled which favor the survival of AOB due to the higher affinity coefficients (Bernet et al., 2001). Higher temperature (30-35 °C) promotes AOB proliferation over NOB due to higher specific growth rates (Hellinga et al., 1998), but in this study, the temperature was constant at 25 °C. pH is closely related to the concentration of FA and FNA. FA (~1 mg/L) and FNA (~9×10^-5^ mg/L) were kept at a level which is much lower than the typical concentration range that was reported to cause inhibition of NOB (1-10 mg/L FA and 0.2-2.8 mcalg/L FNA) (Sinha and Annachhatre, 2007). Overall, those common NOB suppression strategies do not apply in this case. What’s special about this system was the introduction of an anaerobic phase and high concentrations of VFA, therefore, we suspect the extended anaerobic phase or cycling of anaerobic/aerobic phase and high concentration of VFA shocks are the potential selector forces to achieve the partial nitrification. This was further confirmed with independent batch testing with complete nitrifying sludge.

In the extended anaerobic phase, the differential decay rates among different organisms would play a role in population selection. Previous studies reported that NOB had a higher decay rate than AOB (0.306 d^-1^ of NOB and 0.144 d^-1^ of AOB) by measuring Oxygen Uptake Rate (OUR) in a non-fed aerated reactor (Hao et al., 2009). Especially, under the alternating anoxic/aerobic conditions, the difference in decay rates between AOB (0.025 d^-1^) and NOB (0.044 d^-1^) were even more evident (Elawwad et al., 2013). What’s more, NOB was found to exhibit the delay effect during the transition from anoxic to aerobic conditions due to a lag enzyme response, which can contribute to the nitrite accumulation (Kornaros et al., 2010).

On the other hand, a high concentration of VFA shock could be another reason causing nitrite accumulation. Traditionally, rbCOD is at much lower level in the anaerobic or denitrification zones in the biological nutrient removal systems and are less concerned as an important factor. However, for the manure wastewater or side-stream EBPR process (utilizing the return activated sludge to enhance the VFAs production by hydrolysis and fermentation), high concentrations of rbCOD accompany with high VFAs, which can regard as an inhibitor to nitrifiers. Eilersen et al. (1994) reported that acetate acid and propionate acid can inhibit the activity of NOBs but have no impact on AOBs. Delgado et al. (2004) and Oguz (2004) found that under the same concentration of VFAs (~600 mg acetate/L), the inhibition of nitrite oxidation was 1.3 times higher than the ammonia oxidation rate. And they also found that with an initial dose of 150 acetic acid mg/L, nitrite oxidation activity was decreased over 45% but ammonia oxidation activity was only decreased around 20%. Gomez et al. (2000) also mentioned that higher concentration of VFAs can cause more severe inhibition on both ammonia and nitrite oxidation. In our SBR, the concentration of VFAs were 500 mg acetate/L and 280 mg propionate/L, which can cause inhibition on both nitrite and ammonia oxidation and that can explain the pretty low nitrification rate in the system (Table 4). Moreover, AOB are more tolerant of VFAs than NOB, resulting in the nitrite accumulation in the reactor. Furthermore, our nitrification inhibition test further confirmed that the combination of anaerobic condition and high VFAs concentration caused more severe inhibition on NOB activity than on AOB (Table 5) and this inhibition difference will be further magnified after going through periodic anaerobic and high VFA shock in the reactor operation.

### 4.3. Implications

This study demonstrated a potential new strategy to achieve simultaneous high-efficient P removal and stable nitrite accumulation with simpler control. For future work, with the stable nitrite accumulation being achieved in the effluent of EBPR system as demonstrated by this study, part of the influent containing ammonium could be introduced together with the nitrite-dominant effluent, followed by an anammox reactor to achieve simultaneous short-cut N and biological P removal. The wastewater used in this study was synthetic one representing typical digested manure wastewater. It is recognized that the composition of agricultural or other industrial wastewaters are varied and more complicated. The strategy proposed in this study can be verified in future studies with real and wider range of high-strength wastewater such as centrate or various agricultural wastewater for simultaneous phosphorus and short-cut nitrogen removal.

## 5. Conclusions

This study investigated the feasibility of combining EBPR with partial nitrification for treating synthetic digester manure wastewater and the following conclusions can be drawn:

1. EBPR was successfully applied in treating high-strength P wastewaters with the effluent P down to 0.8 ±1.0 mg P/L and the P removal efficiency was 99.5 ±0.8%. *Candidatus* Accumulibacter was the dominant PAO with the relative abundance of 14.2-33.1% and one specific Accumulibacter oligotype was enriched accounting for 64.2-87.9% of the total Accumulibacter abundance. This unique Accumulibacter sub-type was found to be dominant in a full-scale side-stream EBPR system and it is different from those typically detected to be abundant in the conventional EBPR systems.
2. The presence of relatively higher abundance of GAOs (*Defluviicoccus* and *Candidatus* Competibacter) (15.6-40.3%) didn’t deteriorate the EBPR performance.
3. Simultaneous partial nitrification was achieved with the effluent nitrite up to 20.4 ± 6.4 mg N/L and the nitrite accumulation ratio (NAR) was nearly 100% maintained for 140 days (420 cycles). *Nitrosomonas* was the dominant AOB with relative abundance of 0.3-2.4% while NOB was almost undetected (<0.1%).
4. The introduction of extended anaerobic phase and high VFAs concentrations were proposed to be the potential selector forces to promote partial nitrification in combination with EBPR.

## Supplementary Information

E-supplementary data of this work can be found in online version of the paper.

## Acknowledgements

This work was supported by the Water Research Foundation (WRF 4901) and discretionary funding from Cornell University. In addition, one contributing author was funded by the China Scholarship Council (CSC).

