## Supplementary Information for "Combined Enhanced Biological Phosphorus Removal (EBPR) and Nitrite Accumulation for Treating High-strength Wastewater"

### Methodology

#### Oligotyping with 16S amplicon sequencing data

Oligotyping was carried out to further resolve *Candidatus* Accmulibacter genus OTUs into sub-genotypes, namely oligotypes, based on the high-variational sites in reconstructed 16S sequences [1], following the protocol in a previous study [2]. Briefly, five 16S gene V4 region amplicon sequencing datasets (day 32, 61, 111, 144, 158) were analyzed in mothur 1.43.0 [3]. Contigs were aligned to Silva v138 database [4] and classified by MiDAS 3.7 [5], a 16S rRNA database curated specifically for wastewater systems. Only the sequences classified as *Candidatus* Accumulibacter genus were extracted (script:https://github.com/DenefLab/MicrobeMiseq/tree/master/mothur2oligo).

Oligotypes were then curated by resolving the sites with high entropy (Fig. S1), then manually refined until each oligotype contains no high entropy (≥0.2) positions. Five minor oligotypes with less than 41 total read counts were discarded (Fig. S2). The relative abundances of each oligotype were estimated based on read counts. In total 3 oligotypes were identified from the *Candidatus* Accumulibacter genus.

Oligotype 1: CGTTGTGGTCCAAT;

Oligotype 2: CGTTGTGATCCAAT;

Oligotype 3: CGTTGGGGGCCAAT.

**Phylogenetic tree construction**

To reveal the phylogeny of identified *Candidatus* Accumulibacter oligotypes, a phylogenetic analysis was conducted on all identified oligotype representative sequences (3 total), *Candidatus* Accumulibacter oligotypes in a previous study [2] (9 total) and *Candidatus* Accumulibacter phosphatis reference sequences in the MiDAS database [5] (18 total). An extra random sequence (*Dechloromonas*, FLASV96.1460) in the same *Rhodocyclaceae* family was chosen from the MiDAS database as outgroup. Sequences were aligned using MAFFT v7.429 [6]. The phylogenetic tree was then searched using RAxML 8.2.12 [7] and visualized using online tool (https://itol.embl.de/).

**Calculation for nitrogen mass balance for one reactor cycle**

$TN_{in} = TN_{eff} + TN_{deni} + TN_{growth}$

Where:

$TN_{in}$ is the influent TN concentration (mg N/L);

$TN_{eff}$ is the effluent TN concentration (mg N/L);

$TN_{deni}$ is the nitrogen concentration being removed via denitrification (mg N/L);

$TN_{growth}$ is the nitrogen concentration being used for cell growth (mg N/L).

Take the batch cycle on day 132 as an example:

$TN_{in} = 33.12$ mg N/L; $TN_{eff} = 7.02$ mg N/L; $TN_{deni} = 7.90$ mg N/L.

As for $TN_{growth}$, everyday 400 ml mixed liqueur was wasted and based on the measurement, MLSS and MLVSS were kept constant (MLVSS is ~4400 mg/L). Since there were 3 cycles per day, per each cycle 400/3 ml mixed liqueur was wasted which means for each cycle 400/3/1000 L × 4400 mg/L = 586.67 mg biomass was synthesized. The empirical cell biomass formula is $C_5H_7O_2N$.

Therefore, $TN_{growth}$ = (586.67 mg biomass × 14/113)/4L = 18.17 mg N/L.

In this case, $TN_{eff} + TN_{deni} + TN_{growth} = 7.02+7.90+18.17 = 33.09 = TN_{in}$.

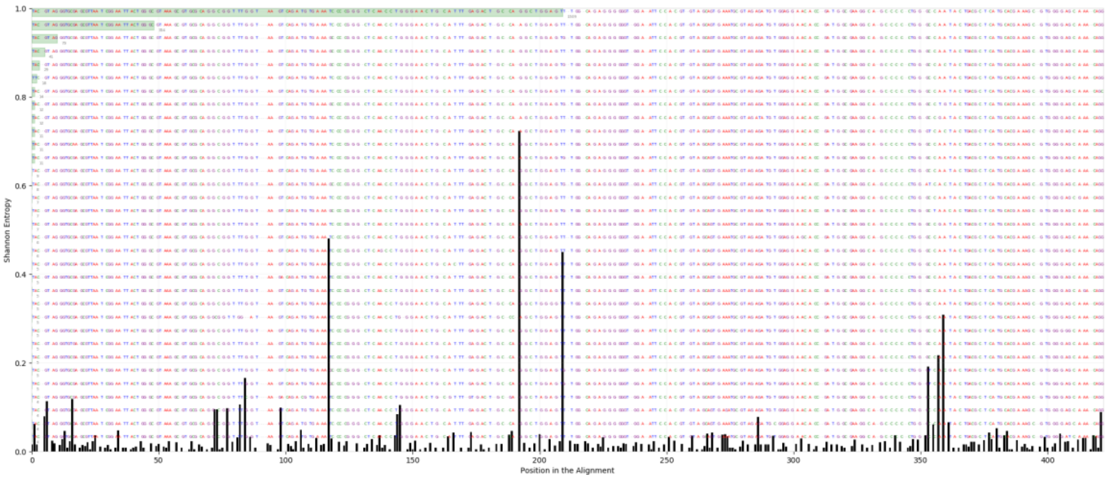


**Fig. S1** Entropy per position calculated in the multiple sequence alignment (MSA) of *Candidatus* Accumulibacter 16S rRNA amplicon sequencing contigs. Higher entropy indicates positions with higher base-type (A, C, G, T, or gap) variations. These positions were used to resolve *Candidatus* Accumulibacter genus OTUs into oligotypes.

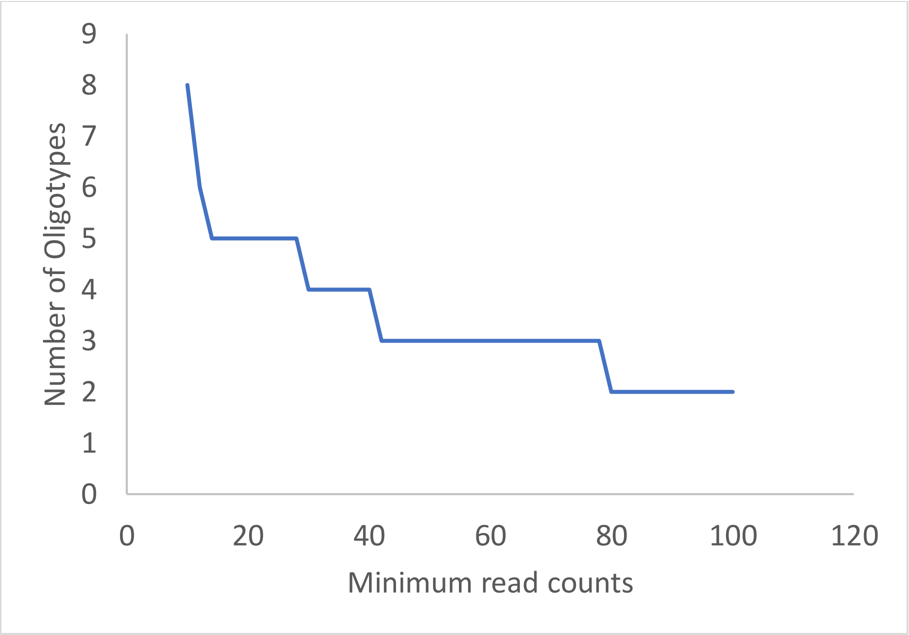


**Fig. S2** Number of oligotypes identified versus minimum total read counts (across all samples) chosen. The initial fast drop of oligotype number (10-40) indicated a large number of minor oligotypes with very low abundances which were considered as noise oligotypes and discarded. The minimal read count parameter was decided to be 41.

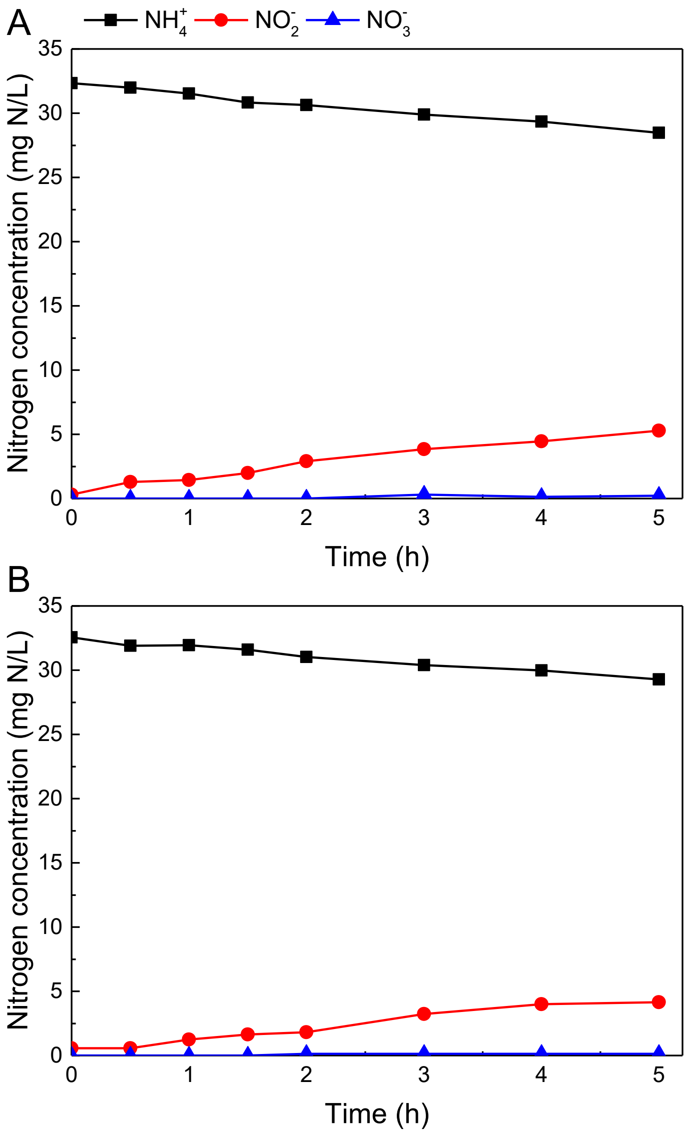


**Fig. S3** Nitrification batch activity test at S2 on day 116.

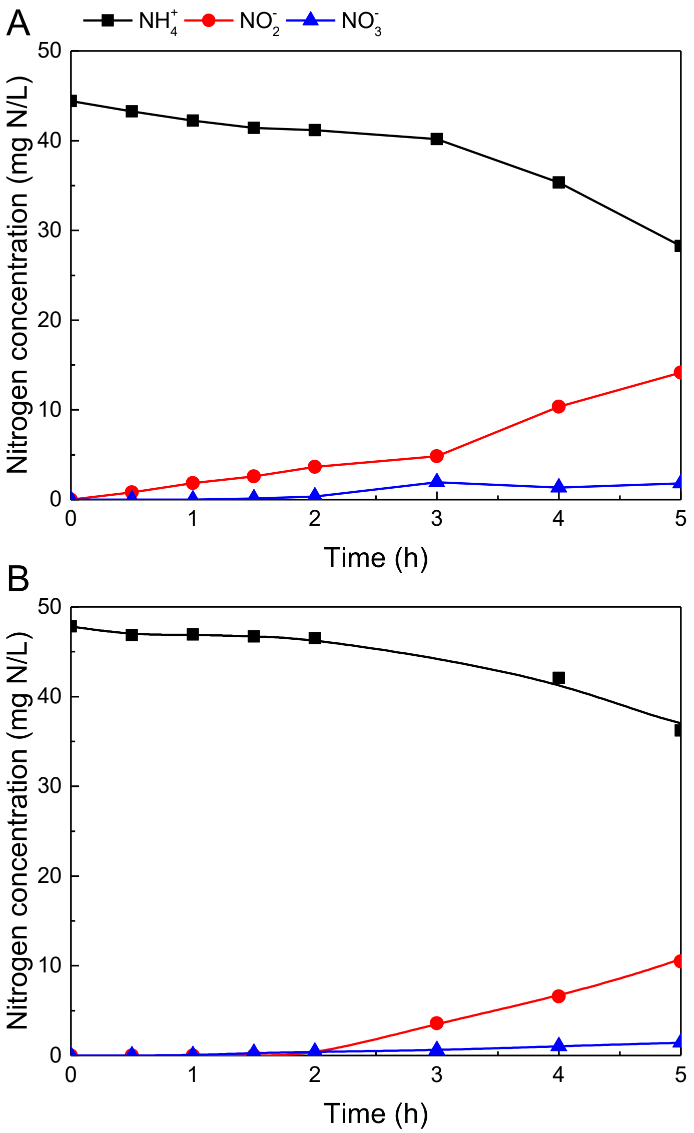


**Fig. S4** Nitrification batch activity test at S3 on day 153.
